# Debiasing FracMinHash and deriving confidence intervals for mutation rates across a wide range of evolutionary distances

**DOI:** 10.1101/2022.01.11.475870

**Authors:** Mahmudur Rahman Hera, N. Tessa Pierce-Ward, David Koslicki

## Abstract

Sketching methods offer computational biologists scalable techniques to analyze data sets that continue to grow in size. MinHash is one such technique to estimate set similarity that has enjoyed recent broad application. However, traditional MinHash has previously been shown to perform poorly when applied to sets of very dissimilar sizes. *FracMinHash* was recently introduced as a modification of MinHash to compensate for this lack of performance when set sizes differ. This approach has been successfully applied to metagenomic taxonomic profiling in the widely used tool `sourmash gather`. While experimental evidence has been encouraging, FracMinHash has not yet been analyzed from a theoretical perspective. In this paper, we perform such an analysis to derive various statistics of FracMinHash, and prove that while FracMinHash is not unbiased (in the sense that its expected value is not equal to the quantity it attempts to estimate), this bias is easily corrected for both the containment and Jaccard index versions. Next, we show how FracMinHash can be used to compute point estimates as well as confidence intervals for evolutionary mutation distance between a pair of sequences by assuming a simple mutation model. We also investigate edge cases where these analyses may fail, to effectively warn the users of FracMinHash indicating the likelihood of such cases. Our analyses show that FracMinHash estimates the containment of a genome in a large metagenome more accurately and more precisely when compared to traditional MinHash, and the point estimates and confidence intervals perform significantly better in estimating mutation distances.

## Introduction

One strategy scientists use when analyzing large data sets is to create a low-dimensional “sketch” or “fingerprint” of their data that allows fast, but approximate answers to their query of interest. Such sketching-based approaches in recent years have been successfully applied to a variety of genomic and metagenomic analysis tasks, due in large part to such methods incurring low computational burden when applied to large data sets. For example, Mash (Ondov et al 2016), is a MinHash (Broder 1997)-based approach that was used to characterize the similarity between all pairs of RefSeq genomes in less than 30 CPU hours. Such efficiency gains are due primarily to sketching-based approaches recording a small subsample (or modification thereof) of the data in such a fashion that some distance metric or similarity measure is approximately preserved, a process called a locality sensitive hashing scheme. In bioinformatics, this has resulted in improvements to error correction (Miclotte et al 2015; Sahlin and Medvedev 2021), assembly (Birol et al 2009; Chin and Khalak 2019; Ekim et al 2021; Ghosh and Kalyanaraman 2017), alignment (Jain et al 2018; Li 2018), clustering (Crusoe et al 2015; Koslicki and Zabeti 2019; Pierce et al 2019; Zhang et al 2014), classification (Breitwieser et al 2018; LaPierre, Alser, et al 2020; LaPierre, Mangul, et al 2019), and so on. Importantly, the accuracy and efficiency of sketching approaches can frequently be characterized explicitly, allowing practitioners to balance between efficiency improvements and accuracy. Often, these theoretical guarantees dictate that certain sketching approaches are well suited only to certain kinds of data. For example, MinHash, which is used in many of the aforementioned applications, has been shown to be particularly well-suited to quantify the similarity of sets of roughly the same size, but falters when sets of very different sizes are compared (Koslicki and Zabeti 2019). This motivated the introduction of the containment MinHash which utilized a MinHash sketch of the smaller set, with an additional probabilistic data structure (a Bloom filter (Bloom 1970)) to store the larger set. While this improved speed and accuracy, this approach can become quite inconvenient for large sets due to requiring a bloom filter to be created for the larger of the two sets.

To ameliorate this, an approach called the “FracMinHash” was recently introduced (Irber et al 2022; Irber Jr 2020) that uses a MinHash hash selection approach but allows sketch size to scale naturally with the size of the underlying data, similar to ModHash dynamic scaling (Broder 1997). These properties allow both Jaccard and containment estimation between FracMinHash sketches, extending the computational advantages of MinHash sketches beyond similar-sized genome comparisons to sequencing datasets of all types. Most notably, FracMinHash enables large-scale metagenome analyses, including genomic and metagenomic similarity assessment, metagenomic taxonomic classification, streaming database searches, and outbreak detection via genomic surveillance (Pierce et al 2019; Viehweger et al 2021). FracMinHash sketching is implemented in a software package called sourmash (Brown and Irber 2016). Independently, and more recently, the same concept of FracMinHash was introduced with the name *universe minimizer* (Ekim et al 2021).

While there is ample computational evidence for the superiority of FracMinHash when compared to the classic MinHash, particularly when comparing sets of different sizes, no theoretical characterization about the accuracy and efficiency of the FracMinHash approach has yet been given. In this manuscript, we address this missing characterization of accuracy and efficiency by deriving a number of theoretical guarantees. In particular, we demonstrate that the FracMinHash approach, as originally introduced, requires a slight modification in order to become an unbiased estimator of the containment index (in terms of expected value). After this, we characterize the statistics of this unbiased estimator and derive an asymptotic normality result for FracMinHash. This in turn allows us to derive confidence intervals and hypothesis tests for this estimator when considering a simple mutation model (which is related to the commonly used Average Nucleotide Identity (ANI) score). We demonstrate the accuracy of our ANI estimates exceeds that of current approaches, particularly at high levels of sequence dissimilarity. We also characterize the likelihood of experiencing an edge case when analyzing real data which allows us to provide a level of confidence along with the estimated containment index. Finally, we support the theoretical results with additional experimental evidence and compare our approach to the frequently used Mash distance (Ondov et al 2016). Many of these theoretical findings have already been implemented into the sourmash (Brown and Irber 2016) computational package (see https://github.com/sourmash-bio/sourmash/pull/1967 and https://github.com/sourmash-bio/sourmash/pull/2032). A Python-based implementation of the theorems we derive is freely available at https://github.com/KoslickiLab/mutation-rate-ci-calculator.

The main part of the manuscript is centered around theoretical analyses using the containment index mainly because of mathematical tractability. The Jaccard index, on the other hand, proved to be more difficult to analyze. We still show how to theoretically analyze the Jaccard index (Section A.7), derive a point estimate of mutation rate (and ANI) using the Jaccard index (Section A.8), and investigate the usefulness of this point estimate in Figure S1. However, because of mathematical intractability, we could not derive a confidence interval around the point estimate, and therefore, decided to include these analyses using the Jaccard index in the supplemental materials. For the sake of continuity from a reader’s perspective, we have included proofs of all theorems in the supplemental materials (Section A.3) as well.

*A note on naming:* As Phil Karlton is reported to have said (Naming things is hard n.d.): “There are only two hard things in Computer Science: cache invalidation and naming things.” The latter certainly holds true in computational biology as well. As noted above, the concept discussed herein has been defined similarly and independently by different authors. Ekim et al. (Ekim et al 2021) referred to the concept as *universe minimizers*, Irber et al. (Brown and Irber 2016; Irber Jr 2020; Pierce et al 2019) called it *scaled MinHash*, and Edgar (Edgar 2021) called them *min code syncmers*. A recent twitter thread (@shenwei356 (Wei Shen) et al. Nov. 21, 2021) involving these authors and others coalesced on the following definitions: *FracHash* is any sketching technique that takes a defined fraction of the hashed elements. As such, Broder’s (Broder 1997) ModHash is one such example of a FracHash. A *FracMinHash* is then a sketch that takes a fraction of the hashed elements, specifically those that hash to a value below some threshold (hence the “min”).

## Methods

### FracMinHash and its statistics

We begin by formally defining a FracMinHash sketch by slightly modifying the definition in (Irber et al 2022). We aim to compare two sequences by computing the containment index from their corresponding FracMinHash sketches (which we refer to using the term *fractional containment index*, and define formally later).

### Definitions and preliminaries

We recall the definition of FracMinHash given in (Irber et al 2022) and reiterate its expected value before extending the statistical analysis of this quantity. Given two arbitrary sets *A* and *B* which are subsets of a domain Ω, the containment index *C*(*A, B*) is defined as 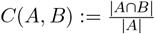 . Let *h* be a perfect hash function *h* : Ω → [0, *H*] for some *H* ∈ ℝ. For a *scale factor s* where 0 ≤ *s* ≤ 1, a FracMinHash sketch of a set *A* is defined as follows:

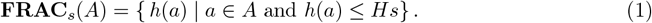

That is, the set of all hashed values in *A* whose hash value is some fraction *s* smaller than the maximum hash value *H*. The scale factor *s* is an easily tunable parameter that can modify the size of the sketch. Using this FracMinHash sketch, we define the FracMinHash estimate of the containment index *Ĉ*_frac_(*A, B*) as follows:

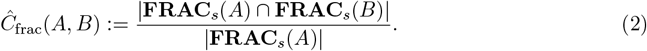

Simply speaking, we want to compute *Ĉ*_frac_(*A, B*) because the sketches are considerably smaller than the original sets *A* and *B*, and we want *Ĉ*_frac_(*A, B*) to accurately approximate *C*(*A, B*). For notational simplicity, let us define *X*_*A*_ := |**FRAC**_*s*_(*A*)|. We observe that if one views *h* as a uniformly distributed random variable, we have that *X*_*A*_ is distributed as a binomial random variable: *X*_*A*_ ∼ Binom(|*A*|, *s*). In practice, hashing libraries use large enough hash value space (i.e. 2^64^) and well enough hash functions that the assumptions on *h* are mostly valid. Furthermore, if *A* ∩ *B* ≠ ∅ where both *A* and *B* are non-empty sets and one is not a subset of the other, then *X*_*A\B*_ and *X*_*A*∩*B*_ are independent when the probability of success, *s*, is strictly smaller than 1. Using these notations, the expectation of *Ĉ*_frac_(*A, B*) is given by Theorem 1, recapitulated from (Irber et al 2022) for completeness.

#### Theorem 1

*For* 0 *< s <* 1, *if A and B are two non-empty sets such that A \ B and A* ∩ *B are non-empty, the following holds:*

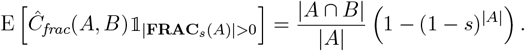

In light of Theorem 1, we note that *Ĉ*_frac_(*A, B*) is *not* an unbiased estimate of *C*(*A, B*): the expected value of *Ĉ*_frac_(*A, B*) is not equal to *C*(*A, B*). This may explain the observations in (Irber Jr 2020) that showed the uncorrected version in eq. (2) leads to suboptimal performance for short sequences (e.g viruses). However, for sufficiently large |*A*| and *s*, the bias factor (1 − (1 − *s*)^|*A*|^) is sufficiently close to 1. Alternatively, if |*A*| is known (or estimated, eg. by using HyperLogLog (Flajolet et al 2007) or the estimate in Section A.6), then

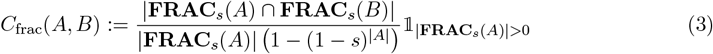

is an unbiased estimate of the containment index *C*(*A, B*). Throughout the rest of the paper, we will refer to the debiased *C*_frac_(*A, B*) as the *fractional containment index*. We now turn to calculating the statistics of *C*_frac_(*A, B*).

### Mean and variance of *C*_frac_(*A, B*)

The expectation of *C*_frac_(*A, B*) is as follows.

#### Theorem 2

*For* 0 *< s <* 1, *if A and B are two distinct sets such that A \ B and A* ∩ *B are non-empty, the expectation of C*_*frac*_(*A, B*) *is given by*

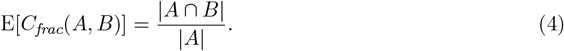

*Proof*. This follows directly from Equation (3) and Theorem 1.

We now turn to determining the variance of *C*_frac_(*A, B*). Ideally, we can do so by using the multivariate probability mass function of *X*_*A*∩*B*_ and *X*_*A\B*_ . However, we found that doing so does not result in a closed-form formula. Therefore, we use Taylor expansions to approximate the variance.

#### Theorem 3

*For n* = |*A* ∩ *B*| *and m* = |*A \ B*| *where both m and n are non-zero, a first order Taylor series approximation gives*

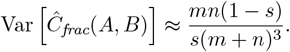

Using the results of Theorem 3 and Equation (3), we have the variance of *C*_frac_(*A, B*) as follows.

#### Corollary 1

*For n* = |*A* ∩ *B*| *and m* = |*A \ B*| *where both m and n are non-zero, a first order Taylor series approximation gives*

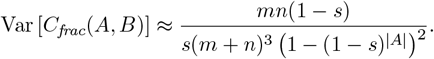

Proceeding in the same fashion, we can obtain series approximations of arbitrarily high order due to the binomial distribution having finite central moments of arbitrary order. However, we found that the higher order expansion derivations are tedious and long, whereas the results obtained using first order approximation are both simple and accurate enough in practice.

### Asymptotic normality of *C*_frac_(*A, B*)

We next prove that *C*_frac_(*A, B*) is asymptotically normal. We utilize the delta method (Agresti 2012) combined with the De Moivre-Laplace theorem, which guarantees asymptotic normality of *X*_*A*∩*B*_ and *X*_*A\B*_, and since 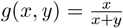 is a function that is twice differentiable, setting *x* = *X*_*A*∩*B*_ and *y* = *X*_*A\B*_ satisfies all requirements of using the delta method on *g*(*x, y*), which gives us the following result:

#### Theorem 4

*For* 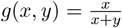, *n* = |*A* ∩ *B*| *and m* = |*A \ B*| *where both m and n are non-zero*,

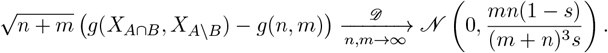

We note that additional statistical quantities can easily be derived. For example, in Section A.5 we provide concentration inequalities that demonstrate theoretically how little *C*_frac_(*A, B*) deviates from its expected value. Section A.6 provides a simple way to calculate the number of distinct *k*mers from a given sketch. Lastly, Section A.7 demonstrates how to compute (and de-bias) a Jaccard estimate from FracMinHash sketches.

### Statistics of *C*_frac_(*A, B*) under simple mutation model

In the previous section, we introduced *C*_frac_(*A, B*), and derived its statistics. In this section, we use these results and connect *C*_frac_(*A, B*) to a biologically meaningful quantity – the Average Nucleotide Identity (ANI) and mutation rate. We do this by assuming a simple mutation model, where each nucleotide of some sequence *S* is independently mutated at a fixed rate, *p*, resulting in the mutated sequence *S*^*′*^ which has expected ANI of 1 − *p* with *S*. This model was recently introduced in (Blanca et al 2022) where it was quantified how this mutation process affects the *k*-mers in *S*.

Before mentioning the details of the mutation model, it is important to note that there are other models of evolution, e.g. TK4 and TK5 models (Takahata and Kimura 1981), the general time reversible (GTR) model (Tavaré 1984) and Sueoka’s model (Sueoka 1995). These vary in the number of parameters used, as well as the degree of complexity. In this work, we consider the simple mutation model because (a) the statistics of *k*-mers under this model are already well explored, and (b) it allows us to connect *C*_frac_(*A, B*) and mutation rate *p* directly, which would not be the case if we considered one of these more nuanced models. The mutation model we use, even though simple enough to be mathematically tractable, is more realistic that the Poisson model assumed by Mash (Ondov et al 2016), which assumes that all *k*-mers are mutated independently, where in reality, one point mutation can affect up to *k* number of *k*-mers. Our experiments reveal that even in case of real genomes, where the lengths of two sequences can be widely dissimilar and clearly the assumptions of the simple mutation model are violated, our approach can accurately determine the mutation rate (and ANI) between two real-world sequences.

We first recall a few important definitions before introducing our findings.

### Preliminaries

Here, we closely follow the exposition contained in (Blanca et al 2022). Let *L >* 0 be a natural number that denotes the number of *k*-mers in some string *S*. A *k-span K*_*i*_ is the range of integers [*i, i* + *k* − 1] which denotes the set of indices of the sequence *S* where a *k*-mer resides. Fix a *mutation rate p* where 0 *< p <* 1. The *simple mutation model* considers each position in *i* = 1, …, *L* + *k* − 1 and with probability *p*, marks it as *mutated*. A mutation at location *i affects* the *k*-spans *K*_max(1,*i*−*k*+1)_, …, *K*_*i*_. Let *N*_mut_ be a random variable defined to be the number of affected/mutated *k*-spans. We use *q* = 1 − (1 − *p*)^*k*^ to express the probability that a *k*-span is mutated. Note that 1 − *p* corresponds precisely to the expected average nucleotide identity (ANI) between a sequence *S* and its mutated counterpart *S*^*′*^.

Given a nonempty sequence *S* on the alphabet {*A, C, T, G*} and a *k*-mer size such that each *k*-mer in *S* is unique, let *A* represent the set of all *k*-mers in *S* and let *L* = |*S*| − *k* + 1. Now, we apply the simple mutation model to *S* via the following: if for any *i* ∈ [1, …, *L* + *k* − 1], this index is marked as mutated, let *S*^*′*^_*i*_ be some nucleotide in {*A, C, T, G*} *\* {*S*_*i*_}, and otherwise let *S*^*′*^_*i*_ = *S*_*i*_. Let *B* represent the set of *k*-mers of *S*, and we assume that *S* does not contain repeated *k*-mers either. In summary, *A* denotes the set of *k*-mers of a sequence *S*, and *B* denotes the set of *k*-mers of a sequence *S*^*′*^ derived from *S* using the simple mutation model with no spurious matches. Note that given a sufficiently large *k*-mer size, these assumptions will be satisfied in most practical scenarios, though the *k*-mer size may be very large (i.e. length of the sequences under consideration). Even so, violations of these assumptions (i.e. repeats and spurious matches) do not negatively impact our results, as demonstrated in the Results section.

We also recall the definition of a confidence interval. Given a distribution and a parameter of interest *τ*, a (1 − *α*)-CI is an interval that contains *τ* with probability 1 − *α*. Given 0 *< α <* 1, we define *z*_*α*_ = *Φ*^−1^(1 − *α/*2), where *Φ*^−1^ is the inverse CDF of the standard Gaussian distribution.

### Expectation and variance of *C*_frac_(*A, B*)

We notice that |*A \ B*| = |*B \ A*| = *N*_mut_, and |*A* ∩ *B*| = *L* − *N*_mut_. We note that the results in Theorem 3, Corollary 1 and Theorem 4 above still hold for a fixed *N*_mut_. However, assuming a simple mutation model, *N*_mut_ is not a fixed quantity, rather a random variable Therefore, the analyses so far only connect *C*_frac_(*A, B*) to a fixed *N*_mut_, as we have only considered the randomness from the FracMinHash sketching process so far. To quantify the impact of the mutation rate *p* on *C*_frac_(*A, B*), we now consider the randomness introduced by both the FracMinHash sketching process and the mutation process simultaneously.

Let 𝒫 = (Ω_1_, *F*_1_, **P**_1_) and 𝒮 = (Ω_2_, ℱ_2_, **P**_2_) be the probability spaces corresponding to the mutation and FracMinHash sketching random processes, respectively. Here, Ω, ℱ and **P** denote the sample space, the sigma-algebra on the sample space, and the probability measure, respectively. We will use the subscript 𝒫, 𝒮 to indicate the product probability space, e.g. E_*𝒫, 𝒮*_ [*·*] and Var_*𝒫, 𝒮*_ [*·*]. Hence we assume that the mutation process and the process of taking a FracMinHash sketch are independent. Before proceeding with the analysis, we make a note that the expectation and variance of *N*_mut_ under the simple mutation model with no spurious matches have been investigated in (Blanca et al 2022). As such, we already know E_*𝒫*_ [*N*_mut_], Var_*𝒫*_ [*N*_mut_] and E_*𝒫*_ [*N*_mut_^2^], and will use these results directly (see Blanca et al 2022, Table 1).

**Table 1:**
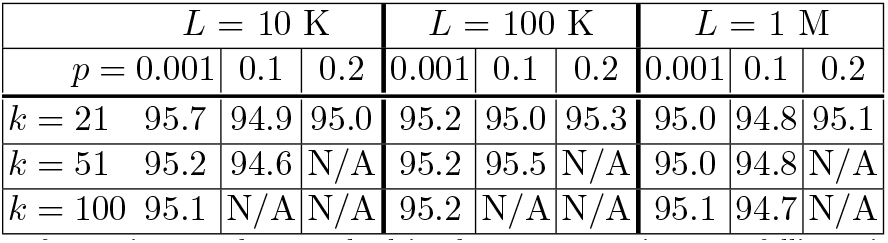
The percentage of experiments that resulted in the true mutation rate falling within the 95% confidence interval given in Theorem 8 when using various mutation rates across multiple *k*-mer sizes and *L* values. A scale factor of *s* = 0.1 was used. The results show an average over 10,000 simulations for each setting. N/A entries indicate that the parameters are not particularly meaningful and will not produce interpretable results, either because *E*[*N*_mut_] ≈ *L* in these cases (almost all *k*-mers are mutated), or because the scale factor is too small to differentiate between the two FracMinHash sketches. These results show that the confidence interval presented in Theorem 8 is statistically significant for meaningful parameter settings. These results reveal that *k*-mers are highly sensitive to mutation rates, and practitioners may need to use shorter *k*-mer lengths to distinguish highly dissimilar sequences.

#### Theorem 5

*For* 0 *< s <* 1, *if A and B are respectively distinct sets of k-mers of a sequence S and a sequence S*^*′*^ *derived from S under the simple mutation model with mutation probability p such that A* ∩ *B is non-empty, then the expectation of C*_*frac*_(*A, B*) *in the product space* 𝒫, 𝒮 *is given by*

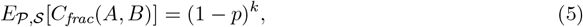

*where* 𝒫 = (Ω_1_, *F*_1_, **P**_1_) *and* 𝒮 = (Ω_2_, *F*_2_, **P**_2_) *are the probability spaces corresponding to the mutation and FracMinHash sketching random processes, respectively*.

To demonstrate how the expected value of *C*_frac_(*A, B*) (considering both the random processes) react to the mutation rate *p* and the *k*-mer size *k*, we show *E*_*𝒫, 𝒮*_ [*C*_frac_(*A, B*)] in a heatmap in Figure 1. The heatmap shows that the expected value of the containment index decreases with a larger mutation rate and a larger *k*-mer size. A point to note is that the expected containment index characterized in Theorem 5 is simply the fraction of non-mutated *k*-mers after undergoing the simple mutation process. Naturally, with larger *k* and/or larger mutation rate *p*, more *k*-mers are mutated, and the expected value of the containment index decreases. This means that in the lighter cells in the heatmap (such as *k* = 20, *p* = 0.001), a small scale factor would suffice – whereas in the darker cells (such as *k* = 100, *p* = 0.1), even a scale factor of 1.0 may not be sufficient because all *k*-mers are mutated. Consequently, a safe and meaningful choice of the scale factor *s* depends on the choices of *p* and *k* – which we discuss in more detail in later in this section.

**Fig. 1:**
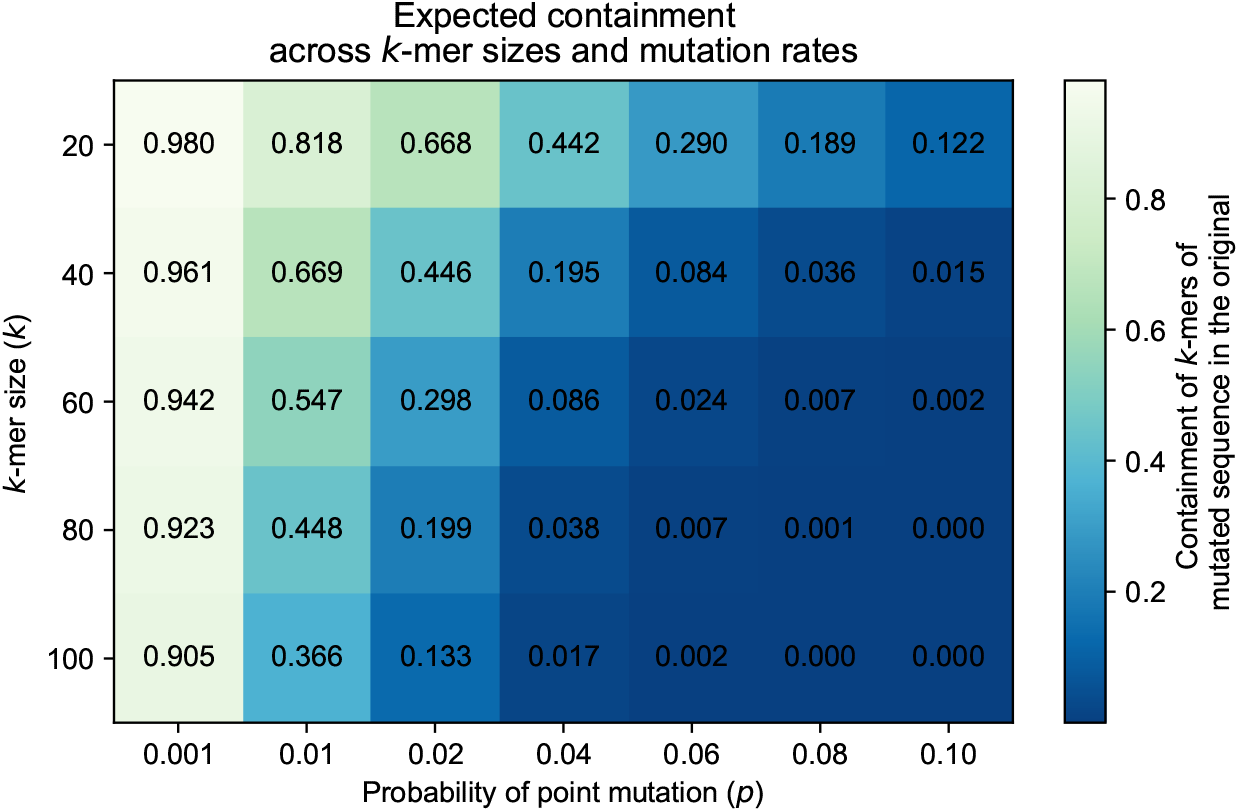
*E*_*𝒫, 𝒮*_ [*C*_frac_(*A, B*)] = (1 − *p*)^*k*^ across different mutation rates and *k*-mer sizes. Darker cells indicate a smaller value. The expected containment decreases with a larger mutation rate, and a larger *k*-mer size. This ideal case indicates that even a scale factor of *s* = 1 will be insufficient for large enough *k* sizes and *p* values.

Next, we turn to the more challenging task of calculating the variance of *C*_frac_(*A, B*) in the product space 𝒫, 𝒮.

#### Theorem 6

*For* 0 *< s <* 1, *if A and B are respectively distinct sets of k-mers of a sequence S and a sequence S*^*′*^ *derived from S under the simple mutation model with mutation probability p such that A* ∩ *B is non-empty, then the variance of C*_*frac*_(*A, B*) *in the product space* 𝒫, 𝒮 *is given by*

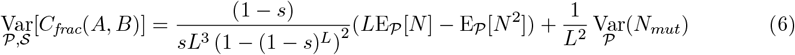

*where* 𝒫 = (Ω_1_, *F*_1_, **P**_1_) *and* 𝒮 = (Ω_2_, ℱ_2_, **P**_2_) *are the probability spaces corresponding to the mutation and FracMinHash sketching random processes, respectively*.

With the results of Theorem 5, we now have a point estimate of the mutation rate *p* given *C*_frac_(*A, B*), which is simply *p* = 1 − *C*_frac_(*A, B*)^1*/k*^ . We next derive a hypothesis test for *C*_frac_(*A, B*) to capture the variability around the point estimate, and later turn it into a confidence interval.

### Hypothesis test and confidence interval

We observe that the marginal of *C*_frac_(*A, B*) with respect to the mutation process is simply 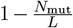 . Using the results in (Blanca et al 2022), we note that *N*_mut_ is asymptotically normally distributed when the mutation rate *p* and *k*-mer length *k* are independent of *L*, and *L* is sufficiently large. In Theorem 4, we showed that *C*_frac_(*A, B*) is normally distributed for a fixed *N*_mut_. Therefore, considering the randomness from both the FracMinHash sketching and the mutation model independently, *C*_frac_(*A, B*) is asymptotically normal when all conditions are met. Using the statistics derived earlier, we obtain the following hypothesis test for *C*_frac_(*A, B*).

#### Theorem 7

*Let* 0 *< s <* 1, *let A and B be two distinct sets of k-mers, respectively of a sequence S and a sequence S*^*′*^ *derived from S under the simple mutation model with mutation probability p, such that A* ∩ *B is non-empty*.

*Also, let* 0 *< α <* 1, *and C*_*low*_ *and C*_*high*_ *be defined as follows*.

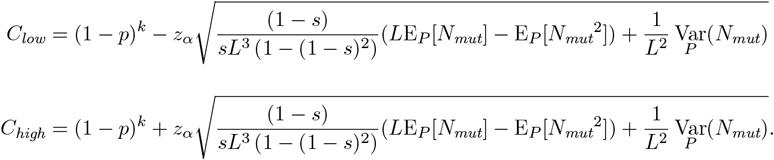

*Then, the following holds as L* → ∞ *and when p and k are independent of L:*

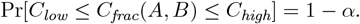

We can turn this hypothesis test into a confidence interval for the mutation rate *p* as follows.

#### Theorem 8

*Let A and B be two distinct sets of k-mers, respectively of a sequence S and a sequence S*^*′*^ *derived from S under the simple mutation model with mutation rate p, such that A* ∩ *B is nonempty. Let* E_*p%fixed*_ [*X*] *and* Var_*p%fixed*_ [*X*] *denote the expectation and variance of a given random variable X under the randomness from the mutation process with fixed mutation rate p*_*fixed*_, *respectively*.

*Then, for fixed α, s, k and an observed C*_*frac*_(*A, B*), *there exists an L large enough such that there exist unique solutions p* = *p*_*low*_ *and p* = *p*_*high*_ *to the following equations, respectively*,

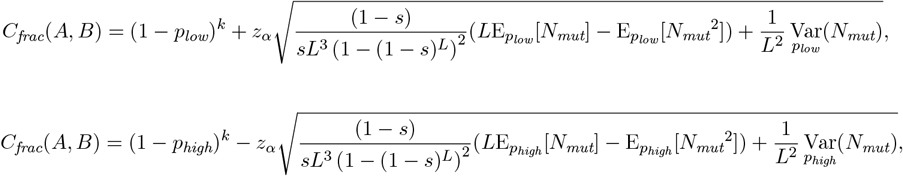

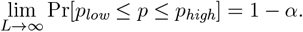

*such that the following holds:*

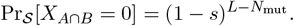

Thus, given an observation of *C*_frac_(*A, B*), along with the point estimate given in Theorem 5, we now have a statistically significant c onfidence interval to lo cate th e mu tation ra te. In th e Results section, we demonstrate that on both simulated and real-world data, the confidence intervals and hypothesis test allow us to accurately connect the mutation rate (and hence, the ANI) with *C*_frac_(*A, B*).

### Setting parameters correctly: likelihood of pathological corner cases

In practice, one disadvantage of sketching techniques is that the size of the sketch (here controlled via the scale factor *s*) may be too small to distinguish between highly similar or dissimilar sequences. For example, given a small mutation rate *p*, one may need a very large scale factor, and so sketch, to be able to distinguish between a sequence and the mutated version. Similarly, if the mutation rate *p* is high and/or a large *k* size is used, it is possible that FracMinHash may report a containment value of 0, even though the true value is nonzero, yet small. These “corner cases” are precisely the ones where the confidence interval given by Theorem 8 will likely fail. One of these pathological cases shows up when there is nothing common between the two FracMinHash sketches **FRAC**_*s*_(*A*) and **FRAC**_*s*_(*B*). We observe that this occurs when *X*_*A*∩*B*_ = 0. Now *X*_*A*∩*B*_ is distributed as a binomial distribution Binom(*n, s*) where *n* = |*A* ∩ *B*| = *L* − *N*_mut_, so the probability of the intersection being empty with respect to the sketching process is:

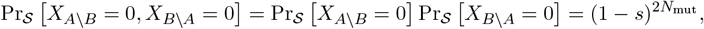

Ideally, we would be able to directly calculate E_𝒫_ [Pr_*S*_ [*X*_*A*∩*B*_ = 0]], the expected probability of this corner case happening. The challenge in doing so is that we do not have a closed form representation of the probability mass function (PMF) of *N*_mut_. As a workaround, we developed a dynamic programming algorithm (presented in Section A.4) to compute Pr[*N*_mut_ = *x*] given the parameters *L* and *p*.

Using this PMF, we can easily compute E_𝒫_ [Pr_*S*_ [*X*_*A*∩*B*_ = 0]], which is the likelihood of the corner case that we observe nothing common between two sequences purely by chance. The remaining pathological case occurs when *p* ≠ 0 and yet **FRAC**_*s*_(*A*) = **FRAC**_*s*_(*B*) (i.e. the sketches are not large enough to distinguish between *A* and *B*). Similar to before, we have

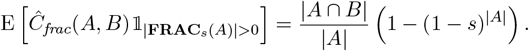

and hence, by calculating E_𝒫_ (1 − *s*)^2*N*mut^ using the PMF of *N*_mut_, we can obtain the likelihood of the latter pathological case. Here, *A \ B* and *B \ A* are disjoint sets, allowing us to use the independence of *X*_*A\B*_ and *X*_*B\A*_. We assume both *A \ B* and *B \ A* are non-empty.

It is important to note the importance to characterize these “corner cases” as without it, a user would be unable to determine if the observed containment index of, say, zero is due to the sequences under consideration being highly diverged, or else the scale factor chosen is much too small. Indeed, these have been implemented (see https://github.com/sourmash-bio/sourmash/pull/ 1860) into sourmash (Brown and Irber 2016) for precisely this purpose – to help practitioners assess if containment estimates of 0 or 1 are due to parameter settings (eg. scale value too high/low), or else are biologically meaningful.

## Results

### FracMinHash accurately estimates containment index for sets of very different sizes

We first show that FracMinHash can estimate the true containment index better than MinHash when the sizes of two sets are dissimilar. For this experiment, we compared FracMinHash with the popular MinHash implementation tool Mash (Ondov et al 2016). We took a *Staphylococcus* genome from the GAGE dataset (Salzberg et al 2012) and selected a subsequence that covers *C*% of the whole genome in terms of number of bases, added this sequence to a metagenome, and calculated the containment of *Staphylococcus* in this “super metagenome.” The metagenome we used is a WGS metagenome sample consisting of approximately 1.3G bases. We used a scale factor of 0.005 for FracMinHash, and we set the number of hash functions for Mash at 4000, since Mash works reasonably well with even only 1000 hash functions to find the containment of *Staphylococcus* genome in the unaltered metagenome. We picked 0.005 because it generates small enough sketch sizes to be computationally inexpensive, and at the same time ensures that the likelihoods of the corner cases are minimal.

We repeated this setup for different values of *C*, and compared the containment index calculated by Mash and FracMinHash in Figure 2. We show the mean values for multiple runs with different seeds in the figure, and use the error bars to show the standard deviation. Mash primarily reports MinHash Jaccard index, so we converted the Jaccard into containment by counting the number of distinct *k*-mers using brute force.

**Fig. 2:**
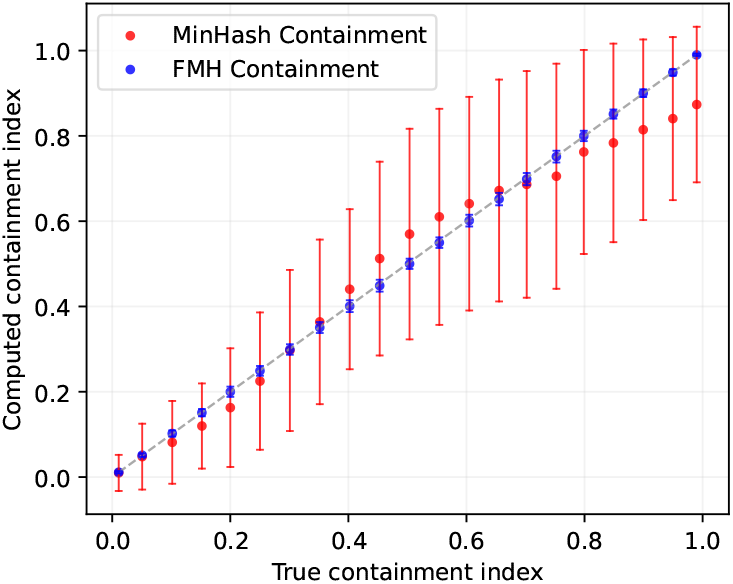
True versus estimated containment index for both Mash and the FracMinHash approach for two sets with dissimilar sizes. The containment index of a *Staphylococcus* genome was computed when *C*% of this genome is inserted into an assembled metagenome. Error bars indicate standard deviation over hash seed values.

Figure 2 illustrates that while Mash and FracMinHash both faithfully estimate the true containment index, the FracMinHash approach more accurately estimates the containment index as this index increases in value. In addition, the estimate is more precise as demonstrated by the size of the error bars on the estimates. This is likely due to the fact that while Mash and FracMinHash both use a sketch of size 4,000 for the *Staphylococcus* genome, Mash uses the same fixed value of 4,000 when forming a sketch for the metagenome, while FracMinHash selects a sketch size that scales with the size of the metagenome. This can be seen most starkly when the metagenome is significantly larger than the query genome.

### FracMinHash gives accurate confidence intervals around mutation rates

Next, we show that the confidence interval from Theorem 8 for the mutation rate *p* is statistically sound and works well in practice. To do so, we performed 10,000 simulations of sequences of length *L* = 10k, 100k and 1M that underwent the simple mutation model with *p* = 0.001, 0.1 and 0.2. We then used a scale factor of *s* = 0.1 when calculating *p*_low_ and *p*_high_ for a 95% confidence interval and repeated this for *k*-mer sizes of 21, 51 and 100. Table 1 records the percentage of experiments that resulted in *p*_low_ ≤ *p* ≤ *p*_high_ and demonstrates that the confidence intervals indeed are approximately 95%.

We also performed the same experiment for other scale factors that also result in minimal likelihood of the corner cases discussed in the Methods section. The results are similar, but for the sake of brevity, these tables are included in the appendix.

In some of these settings (indicated with N/A) shown in Table 1, we skipped the experiment because these do not yield a meaningful result. Mostly, like the darkest cells in Figure 1, these are cases where all or almost all *k*-mers are mutated and consequently, we observe a zero containment, leading to undefined results. In the other cases, the number of shared *k*-mers is too small to use a scale factor of 0.1 and finding a representative number of shared *k*-mers in the FracMinHash sketch – again – resulting in a zero containment. To better understand these settings, we listed the expected number of non-mutated *k*-mers (studied in (Blanca et al 2022)) in Table S3 – only a 10% of which is expected to end up in the sketch, which explains the N/A entries in Table 1.

### FracMinHash more accurately estimates mutation distance On simulated data

We finally compare the Mash estimate and FracMinHash estimate (given as a confidence interval) of mutation rates. For this experiment, we simulated point mutations in the aforementioned *Staphylococcus* genome at a mutation rate *p*, and then calculated the distance of the original *Staphylococcus* genome with this mutated genome using both Mash and the interval given by Theorem 9. The results are shown in Figure 3a. This plot shows that Mash overestimates the mutation rate by a noticeable degree, with increasing inaccuracy as the mutation distance increases. This is likely due to the Mash distance assuming a Poisson model for how mutations affect *k*-mer occurrences, which has been shown to be violated when considering a point mutation model. In contrast, the point estimate given by Theorem 9 is fairly close to the true mutation rate, and the confidence interval accurately entails the true mutation rate.

**Fig. 3:**
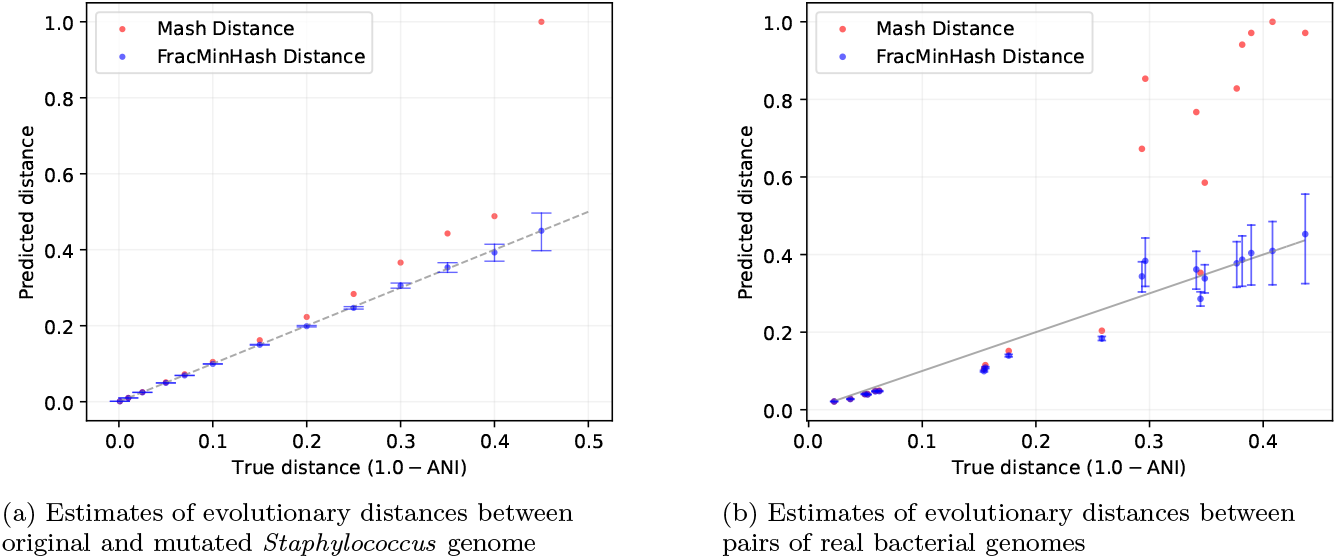
Mash distances and FracMinHash estimates of evolutionary distance (given in terms of one minus the average nucleotide identity: ANI) when (a) introducing point mutations to a *Staphylococcus* genome at a known rate, and (b) between pairs of real bacterial genomes. Error bars indicate the confidence intervals surrounding the FracMinHash estimate calculated using Theorem 8. To obtain the FracMinHash estimates, *k*-mer size *k* = 21 and scale factor *s* = 0.1 was used.

### On real data

We next present pairwise mutation distances between a collection of real genomes using both Mash and the interval in Theorem 8. To make a meaningful comparison, it is important to compute the true mutation distance (or equivalently, the average nucleotide identity) between a pair of genomes. For this purpose, we used OrthoANI (Lee et al 2016), a fast ANI calculation tool. From amongst 199K bacterial genomes downloaded from NCBI, we randomly filtered out pairs of genomes so that the pairwise ANI ranges from 0.5 to 1. For visual clarity, we kept at most 3 pairs of genomes for any ANI interval of width 5%. We used 4000 hash functions to run Mash, and set *L* = (|*A*| + |*B*|)*/*2 for the confidence intervals in Theorem 9, where |*A*| and |*B*| denote the numbers of distinct *k*-mers in the two genomes in a pair. The results are presented in Figure 3b.

Clearly, Mash overestimates the mutation distance, particularly for moderate to high distances. In contrast, the confidence intervals given by Theorem 8 perform significantly better. It is noticeable that the confidence intervals are not as accurate as in the case of a simulated genome (presented in Figure 3a). This is natural because in this real setup, the sizes of the genomes are very dissimilar, have repeats, and very easily violate the simplifying assumptions of the simple mutation model. Nonetheless, these results demonstrate the usefulness of the proposed approach even when the model assumptions are violated.

We conclude this section by computing ANI using FracMinHash sketches (*k* = 21, *s* = 0.1) and a few other well-known tools – namely, Mash (Ondov et al 2016), pyani (Pritchard et al 2016), and fastani (Jain et al 2018). To build the set of genomes for this experiment, we first selected representative genomes from ten species with the largest number of genomes in GTDB-rs207 (7 bacteria and 3 archaea) (Parks et al 2022). Then, we built an “evolutionary path” for each of these “anchor” genomes by selecting three non-representative genomes sharing each taxonomic rank, e.g. three genomes in the same genus but different species, three in the same family but different genus, etc. We computed pairwise ANI from the “anchor” representative genome to each of these genomes using all methods. Using this approach allows usage of real genomes across a range of ANI values. The results are shown in Figure 4. Like the previous set of figures, we again plotted the ANI computed using OrthoANI (Lee et al 2016) on the *x*-axis.

**Fig. 4:**
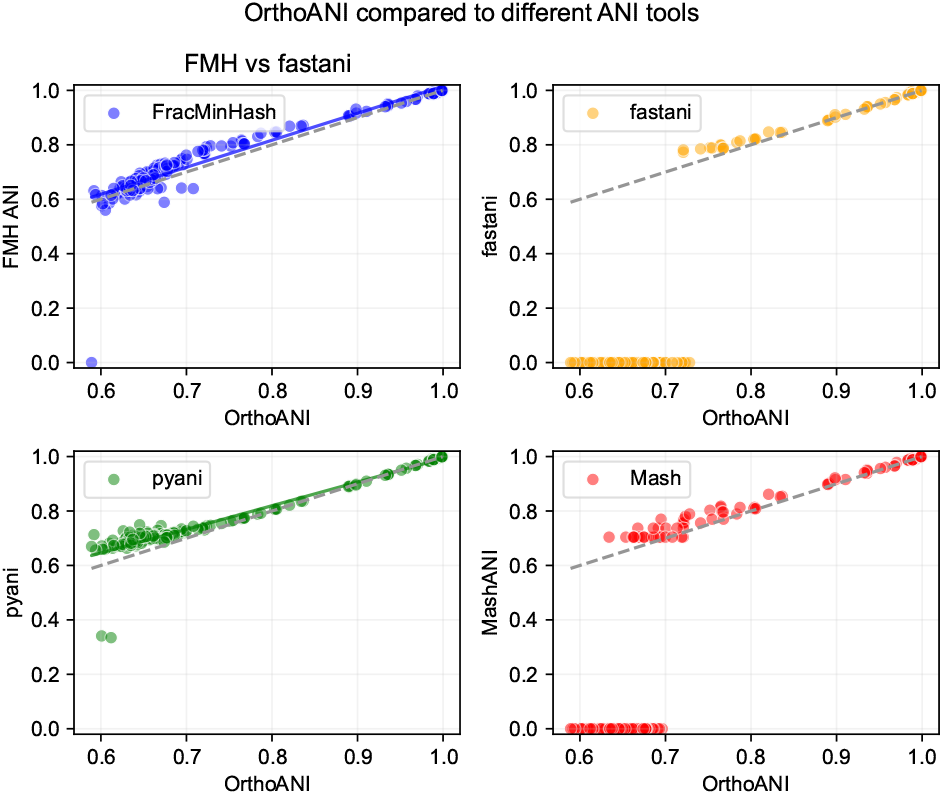
Pairwise ANI estimation among a selection of genomes from the GTDB database. The dashed line represents ANI computed using OrthoANI. The solid blue and green lines show the least-squares fit for the ANI scores computed using FracMinHash and `pyani`, respectively. Mash and `fastani` both have many zero-ed out values, and therefore, the least-squares fit is not shown for these two.

As above, Mash cannot reliably differentiate genomes at ANI below 70%. In addition, as most ANI tools are recommended for use only above 75-80% ANI, the lack of meaningful ANI values below 75% for fastani is expected. In the 80-100% range, all tools correlate well with OrthoANI (shown in dashed grey), with least-squares fits for FracMinHash and pyani in blue and green, respectively. However, both pyani and FracMinHash both correlate with OrthoANI quite well even below 80% ANI, with FracMinHash performing slightly better in this low range. From 60-80% ANI, FracMinHash had a Pearson correlation coefficient with OrthoANI of 0.79 while pyani had a correlation of 0.69 with OrthoANI.

## Discussion

In contrast to classic MinHash, which uses a fixed sketch size, FracMinHash automatically scales the size of the sketch based on the size of the input data. This has the advantage of facilitating accurate comparison of sets of very different sizes, extending sketch-based comparisons to metagenomic datasets, including streaming-based analyses and large-scale database searches. Given that a user has control over what percentage of the data to keep in the sketch (in terms of *s*), reasonable estimates can be made about sketch sizes *a priori*, and trade-offs can be employed to prevent large sketch sizes while maintaining sufficient resolution for search. One particularly attractive feature of FracMinHash is its analytical tractability: as we have demonstrated, it is relatively straightforward to characterize the performance of FracMinHash, derive its statistics, and study how it interacts with a simple mutation model. Given these advantages, it seems reasonable to favor FracMinHash in situations where sets of differing sizes are being compared, or else when fast and accurate estimates of mutation rates are desired (particularly for moderate to high mutation rates). We believe that using FracMinHash can enable fast metagenomic binning by discarding genomes irrelevant to a metagenomic sample (based on low containment scores), let users filter genomes from a reference database using ANI thresholds, allow practitioners to use the confidence intervals with taxonomic tree and sample bootstrap phylogenies off of the taxonomy – and realize many other useful applications.

## Software availability

A Python-based implementation of the algorithms and theorems we derive is freely available at https://github.com/KoslickiLab/mutation-rate-ci-calculator. The results presented in this paper can be reproduced using the code at https://github.com/KoslickiLab/FracMinHash-reproducibles. All code in these github repositories are also available as supplemental materials.

## Acknowledgements

MR and DK were supported by NSF award No. DMS-2029170 and the NIH grant 1R01GM146462. NP was supported by NSF grants 1711984 and 2018911. The authors express their sincere thanks to Luiz Irber and Paul Medvedev for their invaluable inputs to this manuscript.

## Author contributions

DK and MR developed the theoretical results with input from NTP for biological application. MR prepared all the results except for the last figure, the results of which were prepared by NTP. DK and MR wrote the theorems and proofs. MR prepared all plots and the manuscript. All authors reviewed and curated the manuscript.

## A Supplemental materials

### A.1 Verification of Theorem 8 using simulations

Similar to Table 1, we repeated the experiment for the same settings except with two different scale factors. The results are shown in this section.

**Table S1:** The percentage of experiments that resulted in the true mutation rate falling within the 95% confidence interval given in Theorem 8 when using various mutation rates across multiple *k*-mer sizes and *L* values. A scale factor of 0.2 was used. The results show an average over 10,000 simulations for each setting. N/A entries indicate that the parameters are not particularly meaningful and will not produce interpretable results, either because *E*[*N*_mut_] ≈ *L* in these cases (almost all *k*-mers are mutated), or because the scale factor is too small to differentiate between the two FracMinHash sketches.

|  | L = 10 K |  |  | L = 100 K |  |  | L = 1 M |  |  |
| --- | --- | --- | --- | --- | --- | --- | --- | --- | --- |
| p = | 0.001 | 0.1 | 0.2 | 0.001 | 0.1 | 0.2 | 0.001 | 0.1 | 0.2 |
| k = 21 | 95.4 | 95.3 | 94.7 | 95.0 | 95.2 | 95.06 | 95.0 | 95.0 | 94.6 |
| k = 51 | 95.4 | 94.8 | N/A | 94.8 | 94.6 | N/A | 94.9 | 95.1 | 94.4 |
| k = 100 | 94.7 | N/A | N/A | 94.6 | N/A | N/A | 95.4 | 93.7 | N/A |

**Table S2:** The percentage of experiments that resulted in the true mutation rate falling within the 95% confidence interval given in Theorem 8 when using various mutation rates across multiple *k*-mer sizes and *L* values. A scale factor of 0.05 was used. The results show an average over 10,000 simulations for each setting. N/A entries indicate that the parameters are not particularly meaningful and will not produce interpretable results, either because *E*[*N*_mut_] ≈ *L* in these cases (almost all *k*-mers are mutated), or because the scale factor is too small to differentiate between the two FracMinHash sketches.

|  | L = 10 K |  |  | L = 100 K |  |  | L = 1 M |  |  |
| --- | --- | --- | --- | --- | --- | --- | --- | --- | --- |
| p = | 0.001 | 0.1 | 0.2 | 0.001 | 0.1 | 0.2 | 0.001 | 0.1 | 0.2 |
| k = 21 | 96.3 | 95.0 | 96.0 | 95.1 | 95.0 | 95.3 | 95.0 | 95.2 | 94.9 |
| k = 51 | 94.9 | 94.5 | N/A | 94.7 | 95.3 | N/A | 94.7 | 95.0 | N/A |
| k = 100 | 95.2 | N/A | N/A | 95.2 | N/A | N/A | 94.5 | N/A | N/A |

### A.2 Expected number of non-mutated *k*-mers in different scenarios

To explain the N/A entries in Figures 1, S1 and S2, we show the expected number of non-mutated *k*-mers after undergoing the simple mutation process in Table S3. Depending on the scale factor, only a fraction of these non-mutated *k*-mers show up in the FracMinHash sketch. Therefore, if the number of non-mutated *k*-mers is too small, it would be meaningless to run the experiment.

### A.3 Theorems and proofs

#### Theorem 1

**Table S3:**
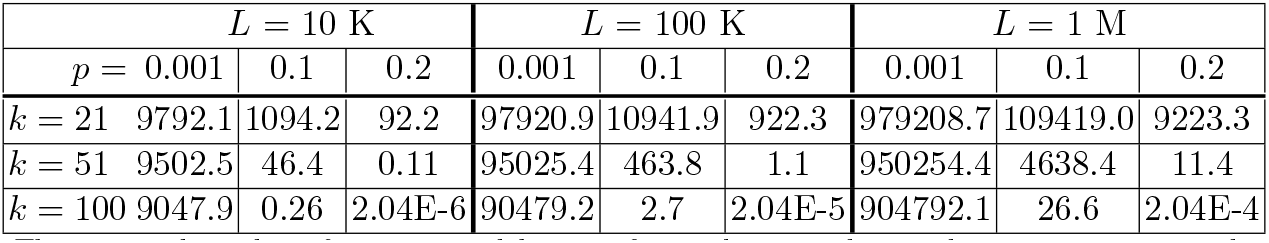
The expected number of non-mutated *k*-mers after undergoing the simple mutation process, shown across multiple *k*-mer sizes, *L* values, and mutation rates.

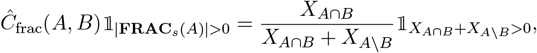

*Proof*. Using the notation introduced previously, observe that

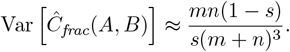

and that the random variables *X*_*A*∩*B*_ and *X*_*A\B*_ are independent (which follows directly from the fact that *A*∩ *B* and *A\ B* are non-empty, distinct sets). We will use the following fact from standard calculus:

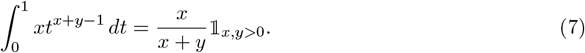

Then using the moment generating function of the binomial distribution, we have

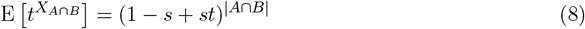

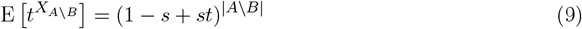

We also know by continuity that

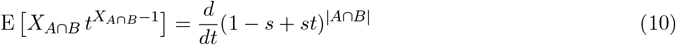

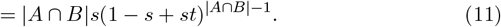

Using these observations, we can then finally calculate that

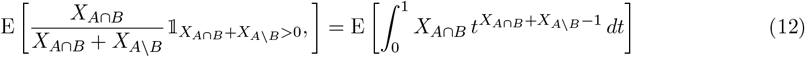

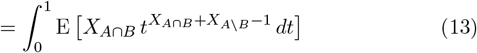

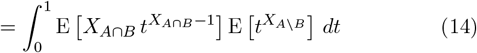

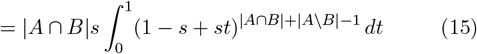

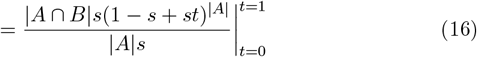

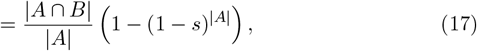

where Fubini’s theorem is used in Equation (13) and independence in Equation (14).

#### Theorem 3

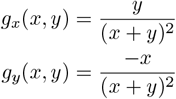

*Proof*. Let *g*(*x, y*) = ^*x*^, *μ*_*x*_ = *ns, μ*_*y*_ = *ms* and use subscripts to denote partial derivatives:

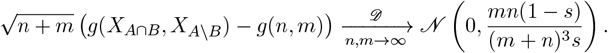

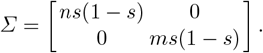

We then have the first order Taylor series:

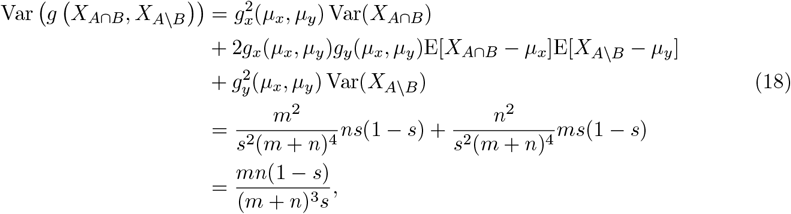

with the middle term of eq. (18) factoring due to independence.

#### Theorem 4

*For* 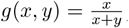,*n* = |*A* ∩ *B*| and *m* = |*A \ B*| *where both m and n are non-zero*,

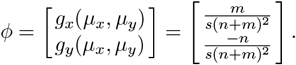

*Proof*. The covariance matrix is calculated as

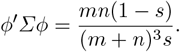

Using the same notation as in Theorem 3, let

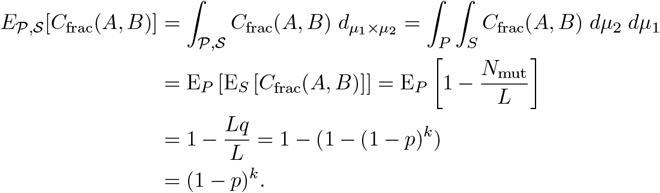

The delta method then uses the first order Taylor series from Theorem 3 to obtain that 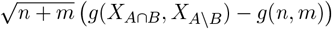 converges in distribution to a centered normal with variance

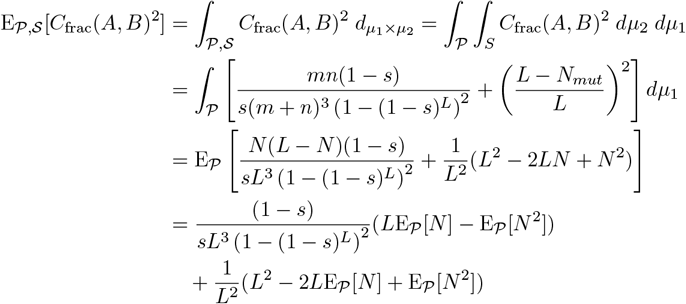

#### Theorem 5

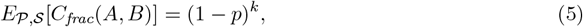

*where* 𝒫 = (Ω_1_, ℱ_1_, **P**_1_) *and* 𝒮 = (Ω_2_, ℱ_2_, **P**_2_) *are the probability spaces corresponding to the mutation and FracMinHash sketching random processes, respectively*.

*Proof*.

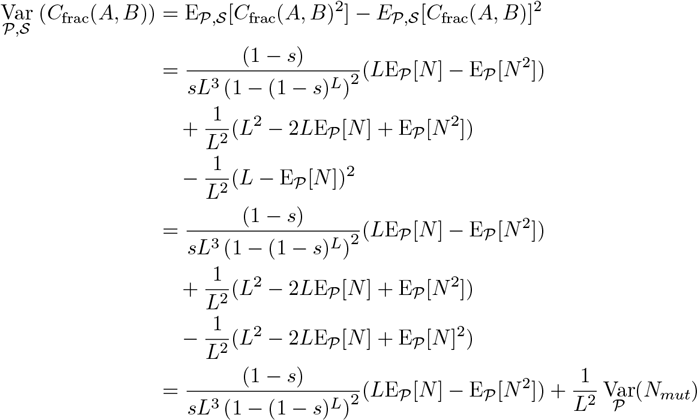

Here, we used Fubini’s theorem in the second step. We also used the expectation of *N*_mut_ from (Blanca et al 2022), where *q* = 1 − (1 − *p*)^*k*^.

#### Theorem 6

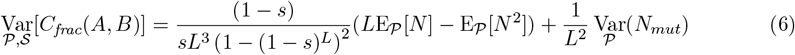

*where* 𝒫 = (Ω_1_, ℱ_1_, **P**_1_) *and* 𝒮 = (Ω_2_, ℱ_2_, **P**_2_) *are the probability spaces corresponding to the mutation and FracMinHash sketching random processes, respectively*.

*Proof*. First, we calculate the second moment of *C*_frac_(*A, B*) in the product space as follows:

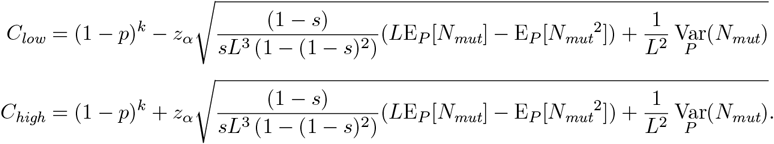

Therefore, we calculate the variance in the product space as follows.

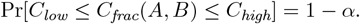

#### Theorem 7

*Also, let* 0 *< α <* 1, *and C*_*low*_ *and C*_*high*_ *be defined as follows*.

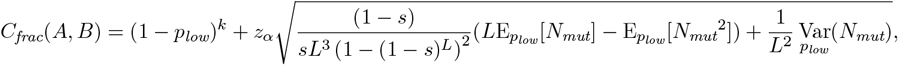

*Then, the following holds as L* → ∞ *and when p and k are independent of L:*

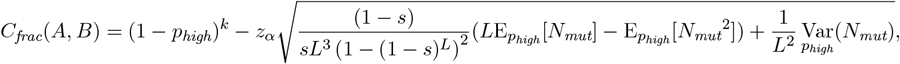

*Proof*. As discussed in the Methods section, *C*_frac_(*A, B*) is asymptotically normal when the required conditions are met. Therefore, the hypothesis test for a random variable following the Gaussian distribution holds for *C*_frac_(*A, B*). Using the expectation and the variance proved in Theorems 5 and 6, we have the results stated in the theorem.

#### Theorem 8

*Let A and B be two distinct sets of k-mers, respectively of a sequence S and a sequence S*^*′*^ *derived from S under the simple mutation model with mutation rate p, such that A* ∩ *B is nonempty. Let* E_*p%fixed*_ [*X*] *and* Var_*p%fixed*_ [*X*] *denote the expectation and variance of a given random variable X under the randomness from the mutation process with fixed mutation rate p*_*fixed*_, *respectively. Then, for fixed α, s, k and an observed C*_*frac*_(*A, B*), *there exists an L large enough such that there exist unique solutions p* = *p*_*low*_ *and p* = *p*_*high*_ *to the following equations, respectively*,

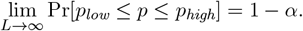

*such that the following holds:*

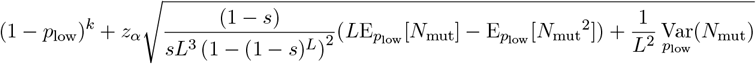

*Proof*. Given the results in Theorem 7, we only need to prove that *p*_*low*_ and *p*_*high*_ are well defined. It suffices to show that

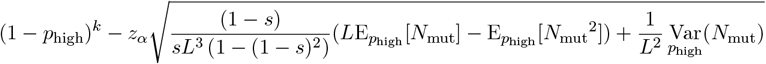

And

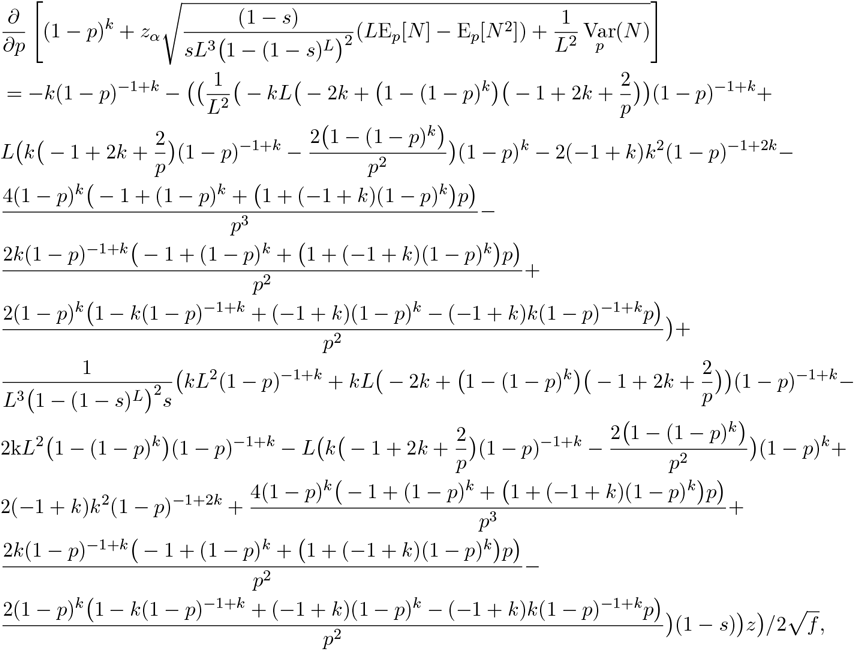

are strictly monotonic in *p*_low_ and *p*_high_, respectively under the Stated conditions.

Let us first investigate the function of *p*_low_. For simplicity, we will write *p* instead of *p*_low_, *z* instead of *z*_*α*_ and *N* instead of *N*_mut_. We observe the following:

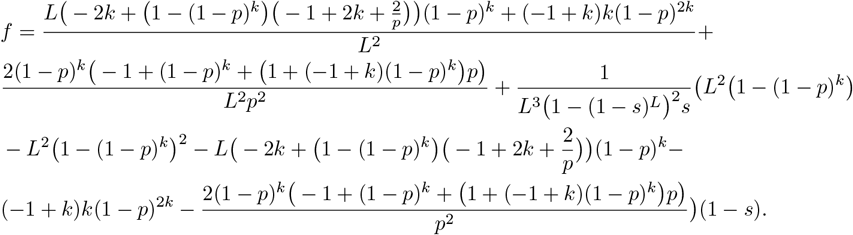

where

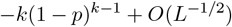

After a tedious, but straightforward (due to the polynomial and rational terms) series expansion of this derivative about *L* = ∞, we obtain that the derivative is

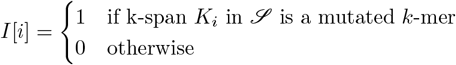

Therefore, as *L* approaches ∞, the derivative is always negative, which gives us that the function 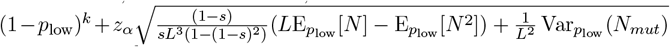 is monotonically decreas-ing in *p*_low_ in the asymptotic case.

The proof that 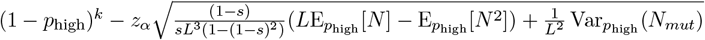 is monotonically decreasing in *p*_high_ proceeds in an entirely analogous manner.

### A.4 Dynamic Programming algorithm to compute the PMF of *N*mut

Here, we will continue to use the notations of the simple mutation model for this algorithm, namely the parameters *L, k* and *p*. Let a string 𝒮 of length *l* undergo the simple mutation process. For ease of understanding, we will represent the mutations introduced to 𝒮 using a binary string ℬ of length *l*, where ℬ [*i*] = 1 if position *i* in 𝒮 was mutated, and 0 otherwise. Therefore, each 1 in this binary string comes from a point mutation, occurring with a probability of *p*, and each 0 with a probability of 1 − *p*. Note that there are *l* − *k* + 1 *k*-mers in 𝒮 . If we could account for all such binary string ℬ’s that result in a total of *x* mutated *k*-mers, we can accumulate the probabilities associated with each of these strings and compute Pr[*N*_mut_ = *x*] by letting *l* = *L* + *k* − 1 (which is the length of *S* and *S*^*′*^). We do this efficiently by defining the following indicator variable:

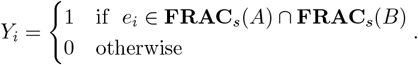

for *i* = 1 up to *l* − *k* + 1, and making use of the following subproblems:

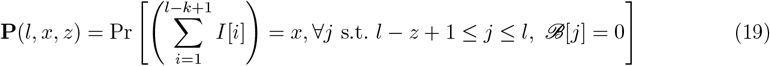

where 0 ≤ *z < k, l* ≥ *k*, 0 ≤ *x* ≤ *l* − *k* + 1. Put another way, **P**(*l, x, z*) is the probability of having *x* mutated *k*-mers in a string of length *l* with *z* trailing zeros. Here, *l* ≥ *k* is required to make sure there is at least one *k*-mer. Equation (19) covers the cases where a string can have at most *k* − 1 trailing zeros. For the rest of the cases, we define the following subproblem:

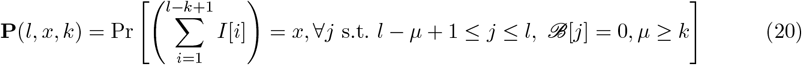

where *l* ≥ *k*, 0 ≤ *x* ≤ *l* − *k* + 1. Put another way, **P**(*l, x, k*) is the probability of having *x* mutated *k*-mers in a string of length *l* with *k* or more trailing zeros.

The base cases of these subproblems are when the string has a length of *k*, and there can only be one *k*-mer. This *k*-mer will be non-mutated when the corresponding binary string has *k* zeros, giving us a probability of **P**(*l* = *k*, 0, *k*) = (1 − *p*)^*k*^. On the other hand, if we have *z < k* trailing zeros, all we need is a 1 preceding these zeros for the *k*-mer to be mutated, giving us **P**(*l* = *k*, 1, *z*) = *p*(1 − *p*)^*z*^ for 0 ≤ *z < k*. It is straightforward to verify that summing these probabilities indeed gives us 1.

We next turn to using the smaller subproblems to solve the larger ones. The core idea is that if we append a 1 at the end of a binary string, then the number of mutated *k*-mers will increase by one, and there are no trailing zeros in the resulting string. On the other hand, if we append a 0 at the end of the string, then the number of mutated *k*-mers will stay the same if the total number of trailing zeros is *k* or more. Appending a 0 at the end will increase the number of mutated *k*-mers by one if the total number of trailing zeros is less than *k*. In both of these latter scenarios, the number of trailing zeros will increase by one. These observations lead to the following recurrence relation:

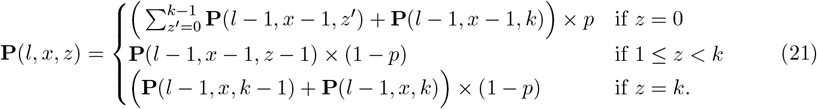

For our parameters *L, k* and *p*, we would need to solve the subproblems for *l* = *L* + *k* − 1. Finally, we would compute Pr[*N*_mut_ = *x*], *x* = 0 up to *L* as follows:

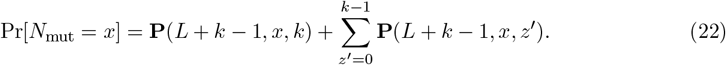

These base cases and recurrence relations give us Algorithm 1 to compute the PMF of *N*_mut_. The loop at Step 5 of the algorithm iterates *L* times. The inner loop at Step 6 iterates at most *L* times. It is straightforward to count that Steps 7 – 11 take *O*(*k*) number of operations. These observations give us a running time of *O*(*L*^2^*k*). Note that *k* is usually in the magnitude of 20 to 50. Considering *k << L*, we have an *O*(*L*^2^) algorithm to compute the PMF of *N*_mut_.

### A.5 Theoretical guarantees to accurately estimate containment index

In this section, we present theoretical evidence that *C*_frac_(*A, B*) is able to estimate the true containment index *C*(*A, B*) with high accuracy. Let the elements in *A* ∪ *B* be *e*_*i*_ for *i* = 1 to *N* . We define an indicator variable *Y*_*i*_ associated with an element *e*_*i*_ as follows:

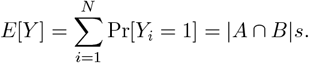

Let *Y* be the number of elements in **FRAC**_*s*_(*A*) ∩ **FRAC**_*s*_(*B*). Naturally, 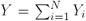. The probability of *Y*_*i*_ being 1 is 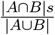 . Therefore, we have:

#### Algorithm 1: PMF − Nmut

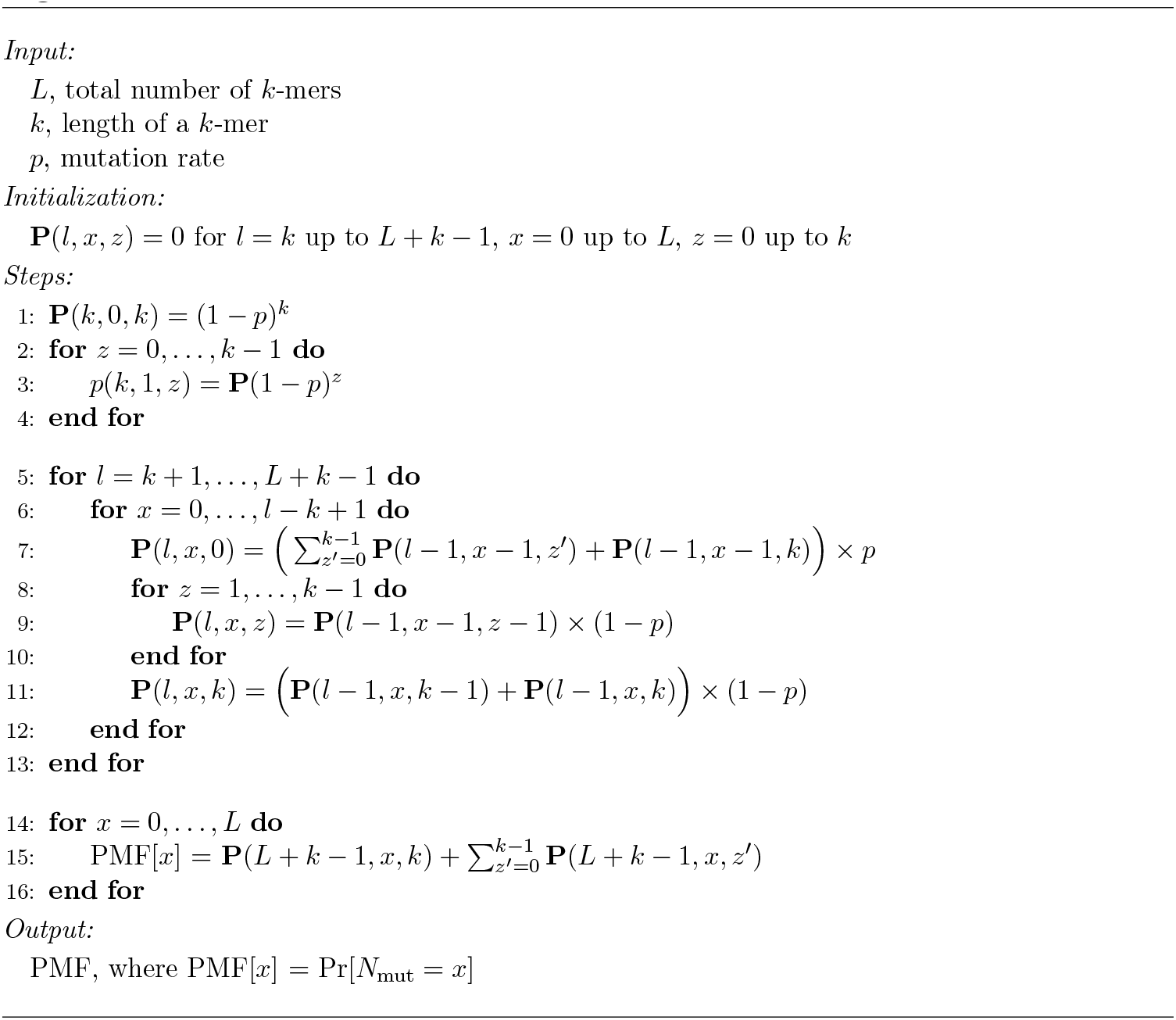

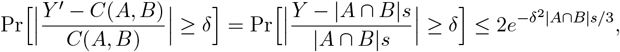

Let us make a simplifying assumption that the exact cardinality of the set *A* is known. Let us define *Y* ^*′*^ as 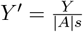. Therefore, *E*[*Y* ^*′*^] = |*A* ∩ *B*|*/*|*A*| = *C*(*A, B*). If we use *Y* ^*′*^ as the estimator to measure *C*(*A, B*), then we have where we used Chernoff bound for a sum of Bernoulli random variables in the last step. The results are trivial, stating that when the two sets have more in common, or when we work with a larger scale factor, the estimate *Y* ^*′*^ performs better. This is expected, and conforms to the concept of using a scale factor. *C*_frac_(*A, B*) estimates *C*(*A, B*) slightly differently than *Y* ^*′*^, and further investigations are required to narrow down the theoretical guarantees of *C*_frac_(*A, B*) estimating *C*(*A, B*).

### A.6 Estimating number of distinct *k*-mers from FracMinHash

In this section, we detail a simple method to estimate the total number of distinct *k*-mers in a given set from its FracMinHash. This can be useful for applications such as the de-biasing in eq. (3) when the set under consideration is small. We have already observed that for *X*_*A*_ := |**FRAC**_*s*_(*A*)| the size of the sketch, *X*_*A*_ is distributed as a binomial random variable: *X*_*A*_ ∼ Binom(|*A*|, *s*). Hence *E*[*X*_*A*_*/s*] = |*A*|, so a point estimate of the number of distinct *k*-mers |*A*| can be had by dividing the sketch size by the scale factor. As the underlying distribution is binomial, we can easily obtain a Chernoff bound for the probability of deviating from this expected value by some relative error *d*:

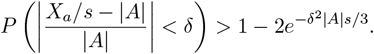

So, for example, if using a scale factor of *s* = 1*/*1000, if one wants to be at least 95% sure that the estimate *X*_*A*_*/s* is off by less than *δ* = 5%, this would require that |*A*| ≥ 4.4 *×* 10^6^.

### A.7 Jaccard calculated using FracMinHash sketches has bias

The theoretical analyses of *C*_frac_(*A, B*) presented in this work reveal the bias in containment index when computed from two FracMinHash sketches. Similarly, a bias in the Jaccard index computed from two FracMinHash sketches can also be proved. Please note that a similar confidence interval can *not* be derived for the Jaccard index as we were able to for the containment index. This is primarily because we found that the Jaccard index cannot be proved to be asymptotically Normal. Nonetheless, the following analysis proves that Jaccard version has a bias associated with it as well.

Let us define

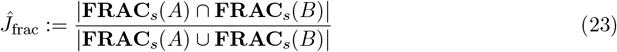

and investigate how well 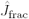 approximates the Jaccard index

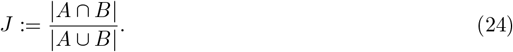

Using the same notations introduced previously, we have the following theorem.

#### Theorem 9

*For* 0 *< s <* 1, *if A and B are two non-empty sets such that A \ B and A* ∩ *B are non-empty and B* ⊄ *A as well as A*⊄ *B, the following holds:*

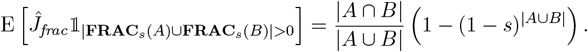

*Proof*. We observe that

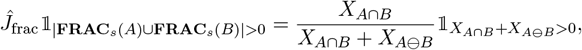

and that the random variables *X*_*A*∩*B*_ and *X*_*A*⊖*B*_ are independent assuming the conditions of the theorem. Here, *A* ⊖ *B* = (*A \ B*) ∪ (*B \ A*). From standard calculus, we have:

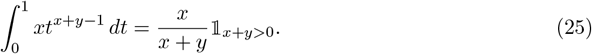

Then using the moment generating function of the binomial distribution, we have

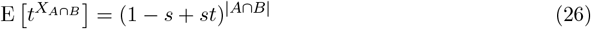

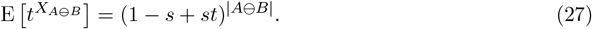

We also know by continuity that

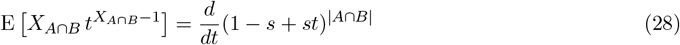

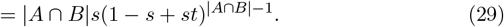

Using these observations, we can then finally calculate that

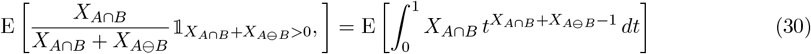

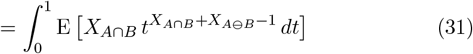

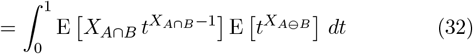

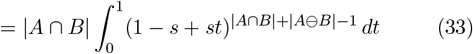

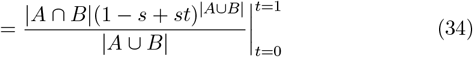

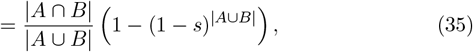

where Fubini’s theorem is used in eq. (31) and independence in eq. (32).

Theorem 9 gives us the following result:

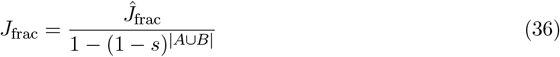

is an unbiased estimator of the Jaccard index *J* . Using the same terminologies introduces in the Methods section, the expectation of this unbiased estimator (considering only the sletching random process) is given by

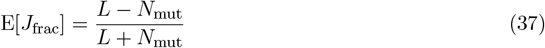

### A.8 Point estimate of mutation rate *p* can be calculated using the Jaccard index

Much like the analysis shown in Theorem 5, we can also obtain a point estimate of the mutation rate *p* using the Jaccard index estimator *J*_frac_.

#### Theorem 10

*For* 0 *< s <* 1, *if A and B are respectively distinct sets of k-mers of a sequence S and a sequence S*^*′*^ *derived from S under the simple mutation model with mutation probability p such that A* ∩ *B is non-empty, then the expectation of J*_*frac*_ *in the product space* 𝒫, 𝒮 *is approximately given by*

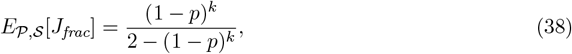

*where* 𝒫 = (Ω_1_, ℱ_1_, **P**_1_) *and* 𝒮 = (Ω_2_, ℱ_2_, **P**_2_) *are the probability spaces corresponding to the mutation and FracMinHash sketching random processes, respectively*.

*Proof*.

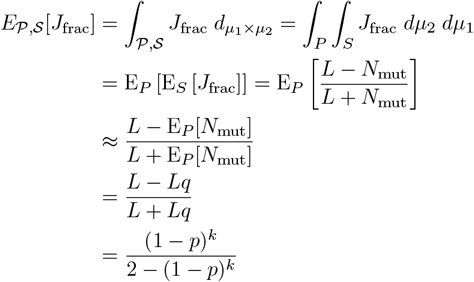

Here, we used Fubini’s theorem in the second step. Using First-Order Taylor approximation, we approximated E[*X/Y*], the expectation of ratio of two random variables, using E[*X*]*/*E[*Y*], the ratio of expectations of the same random variables. We also used the expectation of *N*_mut_ from (Blanca et al 2022), where *q* = 1 − (1 − *p*)^*k*^.

Theorem 10 allows us to obtain the following point estimate of mutation rate *p* using an observed Jaccard index *J*_frac_ obtained through FracMinHash sketches.

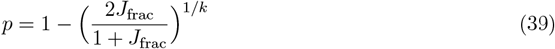

Unfortunately, the random variable 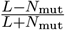 cannot be expressed as Normally distributed –which is the core reason why we could not obtain a confidence interval similar to Theorem 8 using the Jaccard index.

Nevertheless, the point estimate shown above can still be useful. To demonstrate the usefulness of the point estimate obtained in Equation (39), we ran the same set of experiments presented in Figure 3a and Figure 3b. The results are shown in Figure S1 – which show that the point estimate obtained using the Jaccard index is pretty close to the point estimate obtained using the containment index, both in the cases of simulated and real data.

**Fig. S1:**
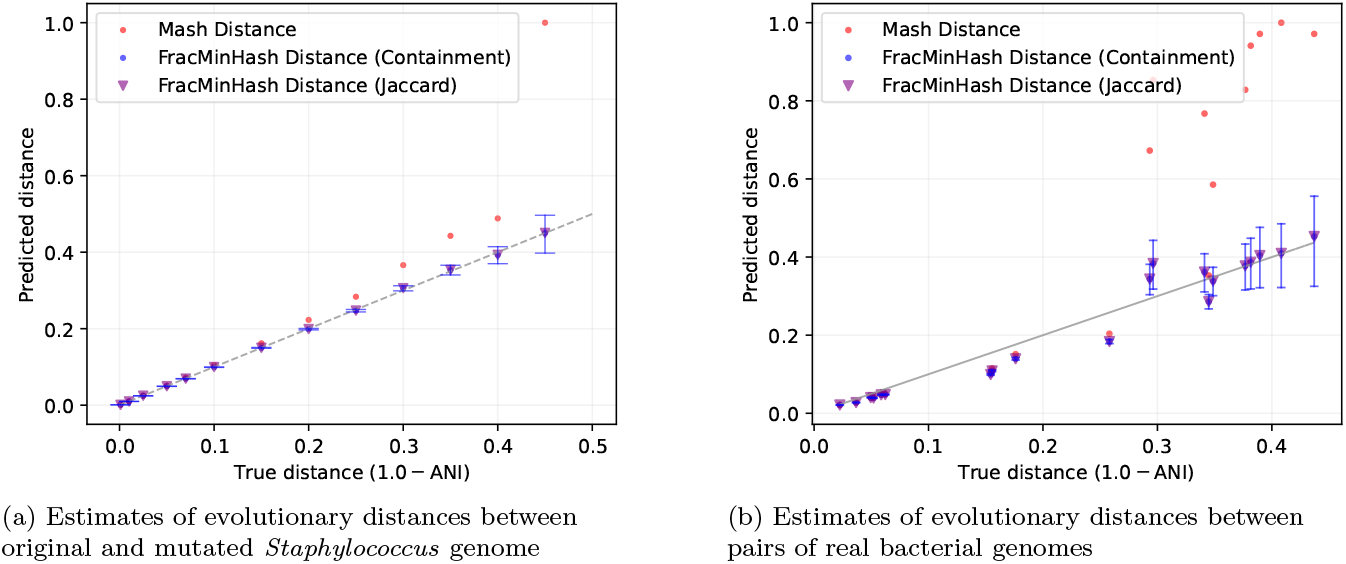
Mash distances and FracMinHash estimates of evolutionary distance (given in terms of one minus the average nucleotide identity: ANI) when (a) introducing point mutations to a *Staphylococcus* genome at a known rate, and (b) between pairs of real bacterial genomes.

